# Prediction of Q-CHAT scores based on Functional Connectivity in Healthy Newborns

**DOI:** 10.64898/2026.06.30.735645

**Authors:** Mi Zou, Arun L. W. Bokde

## Abstract

Neonatal resting-state functional connectivity may provide early markers of later variation in Q-CHAT scores, measured dimensionally within a non-clinical population, but the large-scale systems carrying the most robust predictive signal remain unclear. Using resting-state fMRI data from 397 infants in the Developing Human Connectome Project (277 term-born, 120 preterm-born), we applied a stability-driven, ROI-constrained connectome-based predictive modeling framework to predict 18-month Quantitative Checklist for Autism in Toddlers (Q-CHAT) scores. Significant prediction was observed in the whole cohort and in term-born infants, but did not reach statistical significance in the preterm-only group. Across the statistically significant models (whole cohort and term-born infants), the most prominent hubs were located in occipital and adjacent cortical regions, including the middle occipital gyrus, lingual gyrus, calcarine gyrus, and rolandic operculum. At the network level, the strongest predictive connections linked visual and visual-association systems with auditory networks, with additional contributions from medial motor, temporoparietal, and prefrontal systems. These findings suggest that later variation in Q-CHAT scores, is associated with neonatal large-scale functional organization, particularly in sensory and multisensory pathways.

## Introduction

From the earliest weeks of life, the neonatal brain already displays organized large-scale functional networks, and primary sensory and motor systems can be identified by late gestation and at term-equivalent age (Doria et al., 2010; Fransson et al., 2009). This early architecture provides a scaffold for later development and makes resting-state fMRI (rs-fMRI) a promising tool for identifying early biomarkers of neurodevelopmental variation (Zhang et al., 2019).

Preterm birth, affecting about 11% of live births worldwide, occurs during a period of rapid brain maturation and is associated with altered perinatal connectivity and elevated neurodevelopmental risk (Chawanpaiboon et al., 2019; Eyre et al., 2021; Rogers et al., 2018). Preterm birth is associated with an increased likelihood of atypical neurodevelopmental outcomes (Rogers et al., 2018). An important question, therefore, is whether neonatal functional organization is associated not only with broad developmental outcomes, but also with the early emergence of autism-related traits, and how this association may be altered by preterm birth.

Behavioural traits associated with autism develop gradually across infancy and toddlerhood rather than appearing abruptly. Increasing evidence suggests that their neural antecedents may be detectable before overt behaviours are evident (Girault & Piven, 2020). Prospective imaging studies have implicated atypical sensory and social brain development, with particular emphasis on visual pathways and their relation to later autism-related features (Girault et al., 2022). Recent neonatal work further showed that dynamic functional connectivity at birth is associated with later Q-CHAT variation, including atypical social, sensory, and repetitive behaviours (França et al., 2024).

The Q-CHAT was designed to measure autism-related traits as a quantitative, continuously distributed dimension rather than as a categorical diagnosis, and such traits are approximately normally distributed across the general population, with diagnosed autism representing one tail of a continuum rather than a discrete category (Allison et al., 2012; Constantino & Todd, 2003). In an unselected, predominantly healthy cohort such as the present one, Q-CHAT scores therefore index normal individual variation in early social attention, sensory responsiveness, and communicative behaviour. This variation is informative even when scores lie within the typical range: it is heritable, reasonably stable over time, and continuous with the traits that are elevated in clinical autism, so mapping its neural correlates speaks both to normative neurodevelopment and to the early organisation of systems implicated in autism. Accordingly, the aim here is to model this continuous trait variation and its neonatal connectomic substrate, not to predict or diagnose autism in individual neonates.

Autism is highly heritable, but genetic liability is not expressed directly as behaviour; it acts through the development and organisation of brain circuits, which can be measured with imaging well before autism-related behaviours emerge (Girault & Piven, 2020; Girault et al., 2022). Resting-state fMRI thus provides an intermediate neural phenotype linking inherited liability to later behavioural variation, and the neonatal period offers a window in which this early, partly heritable brain organisation can be observed with minimal influence from postnatal experience. Demonstrating that neonatal functional connectivity carries reproducible, out-of-sample information about later Q-CHAT variation is therefore a necessary step toward using early imaging to understand how genetic and perinatal factors shape the developmental trajectory of autism-related traits.

Despite these advances, three limitations remain. First, many neonatal studies focus on general neurodevelopment or predefined circuits rather than autism-related traits modelled as a continuous, dimensional measure such as that provided by the Q-CHAT. Second, most prior work is association-based rather than predictive. Third, it remains unclear which large-scale systems carry the most stable and interpretable signal for later autism-related variation, and whether the neural networks differ between term-born and preterm-born infants.

These issues are especially relevant in prematurity. Preterm birth is associated with altered structural and functional connectivity, which may disrupt the integration of sensory and higher-order systems during a critical developmental window (Eyre et al., 2021; Rogers et al., 2018). Because early social communication depends on coordinated sensory and associative processing, variation in autism-related traits may be particularly sensitive to such early connectomic differences.

Here, using neonatal rs-fMRI data from 402 infants in the Developing Human Connectome Project, we applied a region-of-interest (ROI)-constrained variant of Connectome-Based Predictive Modeling (CPM) to predict 18-month Q-CHAT scores. CPM is a data-driven framework for linking whole-brain connectivity to behavioral variation and supporting out-of-sample prediction (Shen et al., 2017). Our implementation extended standard CPM with a stability-driven, hub-oriented feature selection strategy.

This approach was motivated by the idea that predictive performance in neonatal rs-fMRI may be weakened by unstable or low-SNR edges. Functional hubs are highly connected and centrally positioned within the brain network, making them plausible anchors for robust predictive signal (Power et al., 2013; van den Heuvel & Sporns, 2013). By identifying stable high-degree ROIs and constraining feature selection to edges connected to them, we aimed to improve both interpretability and reliability in a dataset characterized by substantial developmental variability and motion-related confounds (Fitzgibbon et al., 2020).

We expected Q-CHAT-related prediction to involve distributed sensory and higher-order systems rather than a single circuit. In particular, we anticipated contributions from visual, auditory, temporoparietal, and prefrontal networks, given their roles in sensory integration, social orienting, and communicative processing (Girault et al., 2022; Michon et al., 2022; França et al., 2024). We also examined the whole cohort and birth-status subgroups to test whether predictive organization differed between term-born and preterm-born infants.

## Methods

### Participants

We analyzed data from 397 participants selected from the 814 available functional MRI (fMRI) datasets in the Developing Human Connectome Project(dHCP, Release 4). Among 124 preterm-born infants, 91 had two imaging sessions; in such cases, only the first session, scanned shortly after birth, was included in the analysis. We also performed a supplementary analysis using the term-equivalent-age sessions for comparison. Participants were excluded based on the following criteria: (i) excessive head motion (n = 151; motion exclusion criteria see below), (ii) missing Q-CHAT scores at 18 months (n = 121), and (iii) term-born infants with postnatal age (scan age - gestational age) >3 weeks (n = 54). The final cohort comprised 277 term-born and 120 preterm-born infants, with preterm defined as gestational age <37 weeks. Demographic information is presented in Table 1. Neurodevelopmental outcomes were assessed at 18 months’ corrected age using the Quantitative Checklist for Autism in Toddlers (Q-CHAT).

**Table 1.** Research participants.

|  | Term(n=277) | Preterm(n=120) | Preterm(n=120) | Statistics | *p* value |
| --- | --- | --- | --- | --- | --- |
|  |  |  | term-equivalent age |  |  |
| Demographics |  |  |  |  |  |
| GA at birth,<br>median(range),weeks | 40.14 (37.0-42.7) | 33.29 (23.0-36.9) | 33.29 (23.0-36.9) | 15.8<sup>a</sup> | p < 0.001 |
| PMA at scan,<br>median(range),weeks | 41.14 (37.4-44.7) | 35.41 (27.3-45.1) | 39.67(37.2-42.8) | 13.5<sup>a</sup> | p < 0.001 |
| Female(%) | 125 (45) | 51 (42) | 51 (42) | 0.2<sup>b</sup> | 0.629 |
| Weight at birth,<br>median(range),kg | 3.77 (2.7-4.3) | 1.64 (0.5-4.1) | 1.64 (0.5-4.1) | 10.8<sup>a</sup> | p < 0.001 |
| % FD outliers(SD) | 4.25 (2.7) | 4.75 (2.4) | 4.75 (2.4) | -0.5<sup>a</sup> | p = 0.650 |
| Follow-up |  |  |  |  |  |
| Q-CHAT score,<br>mean(SD)<sup>c</sup> | 30.43 (8.9) | 30.13 (10.3) | 30.13 (10.3) | 0.7<sup>a</sup> | p = 0.496 |

FD framewise displacement
<sup>a</sup>Z(Mann-Whiteney U test)
<sup>b</sup> $\chi^2$ -test
<sup>c</sup>Quantitative Checklist for Autism in Toddlers
The descriptive statistics compare the preterm cohort of the first session and the term cohort.

### MRI Data Acquisition

Functional MRI data were collected as part of the developing Human Connectome Project (dHCP) at the Evelina Newborn Imaging Centre, Evelina London Children’s Hospital, utilizing a 3 Tesla Philips Achieva system. The study received ethical clearance from the UK National Research Ethics Authority (14/LO/1169), and all participating families provided written informed consent prior to the imaging sessions. All scans were performed without the use of sedation in a neonatal environment designed specifically for safety and comfort, which included a custom 32-channel head coil, an acoustic hood, and devices to ensure proper positioning of the infants. Infants wore MRI-compatible ear protection to reduce noise exposure, and a neonatal nurse or paediatrician continuously monitored vital signs, including heart rate, oxygen saturation, and body temperature.

Blood-oxygen-level-dependent (BOLD) fMRI data were acquired using a multi-slice echo planar imaging sequence with multiband excitation (factor 9) over a scan duration of 15 minutes and 3 seconds, producing 2300 volumes. Imaging parameters included a repetition time (TR) of 392 ms, echo time (TE) of 38 ms, a flip angle of 34°, and a voxel resolution of 2.15 mm isotropic. High-resolution T2w anatomical images were obtained for structural analysis and functional data registration. T2-weighted images were obtained with a resolution of 0.8 mm isotropic and a field of view of 145 × 145 × 108 mm, a TR of 12 s, and a TE of 156 ms.

### Functional data preprocessing

The preprocessing of neuroimaging data was performed using a bespoke pipeline specifically optimized for neonatal imaging and developed for the dHCP, as detailed in Fitzgibbon et al. (2020). The pipeline accounted for susceptibility-induced dynamic distortions as well as intra-and inter-volume motion artifacts. Twenty-four extended rigid-body motion parameters were regressed alongside single-subject independent component analysis (ICA) noise components identified using the FSL FIX tool (Oxford Centre for Functional Magnetic Resonance Imaging of the Brain’s Software Library, version 5.0). The denoised data were first registered to the T2-weighted native space using boundary-based registration. They were then non-linearly transformed to a standard space with a weekly template from the dHCP volumetric atlas through diffeomorphic multimodal (T1/T2) registration.

Because head motion is a potential surrogate marker for the infant’s arousal state and can interact with underlying neural activity, we adopted a conservative approach to minimize the impact of motion-related artifacts. Specifically, for each subject, we selected a continuous subset (1600 from the original 2300 acquired volumes) with the minimum total framewise displacement (FD) and the dataset was cropped accordingly – the cropped subset was used for all subsequent analyses. After the cropping, volumes were flagged as motion outliers based on FD, with outliers identified as FD >1.5 interquartile range (IQR) above the 75th centile. Subjects with more than 160 motion-outlier volumes (>10% of data) were excluded from further analyses. The motion outliers of the resulting data from the participants are showed in Table 1. No significant differences were observed between term and preterm groups under the assumption of non-normality (p = 0.650, Mann-Whitney U-test). The number of outliers was included as a covariate in subsequent regression analyses.

### Connectivity matrices

Whole-brain functional connectivity was computed using the CONN toolbox. Brain regions were defined using the Schaefer 200-node cortical atlas combined with 8 subcortical regions from the dHCP parcellation, yielding 208 nodes. The atlas was aligned to the 40-week template via rigid-body, affine, and SyN diffeomorphic transformations implemented in ANTs. Mean time courses were extracted for each ROI, and pairwise Pearson correlations were computed and Fisher Z-transformed, resulting in a symmetric 208×208 connectivity matrix per subject.

### ROI-constrained Connectome–Based Predictive Modeling (R-CPM)

The pipeline was applied separately to preterm-only, term-only, and whole cohort (term + preterm), with preterm status included as a covariate in the whole cohort analysis. All analyses are performed separately for positive and negative associations.

Stage 1 — Stable edge discovery: We ran 10-fold repeated 150 times with new partitions. For each fold we computed partial correlations between each edge and behaviour controlling for gender, head motion (number of outliers), age at scan, and preterm status when applicable. Edges with r>0 and p<0.05 were counted as positive; r<0 and p<0.05 as negative. Counts were accumulated across all folds and repeats. An edge was labelled stable if selected in ≥95% of the 10×150 iterations. ROI ranking(below) was performed separately for positive and negative graphs.

Stage 2 — ROI ranking: From the Stage-1 stable graph (sign-specific), we identified hubs by iteratively removing the highest-degree ROI and recomputing degrees on the remaining graph. The ROI with the largest degree was labelled the top hub; after removing it and its incident edges, the ROI with the largest updated degree was labelled second, and so on, yielding a ROI order *R= (r_1_, r_2_,…)* (Figure 1a). No Stage-1 edges were carried forward as features; only the ROI order *R* was passed to Stage 3.

**Figure 1.**
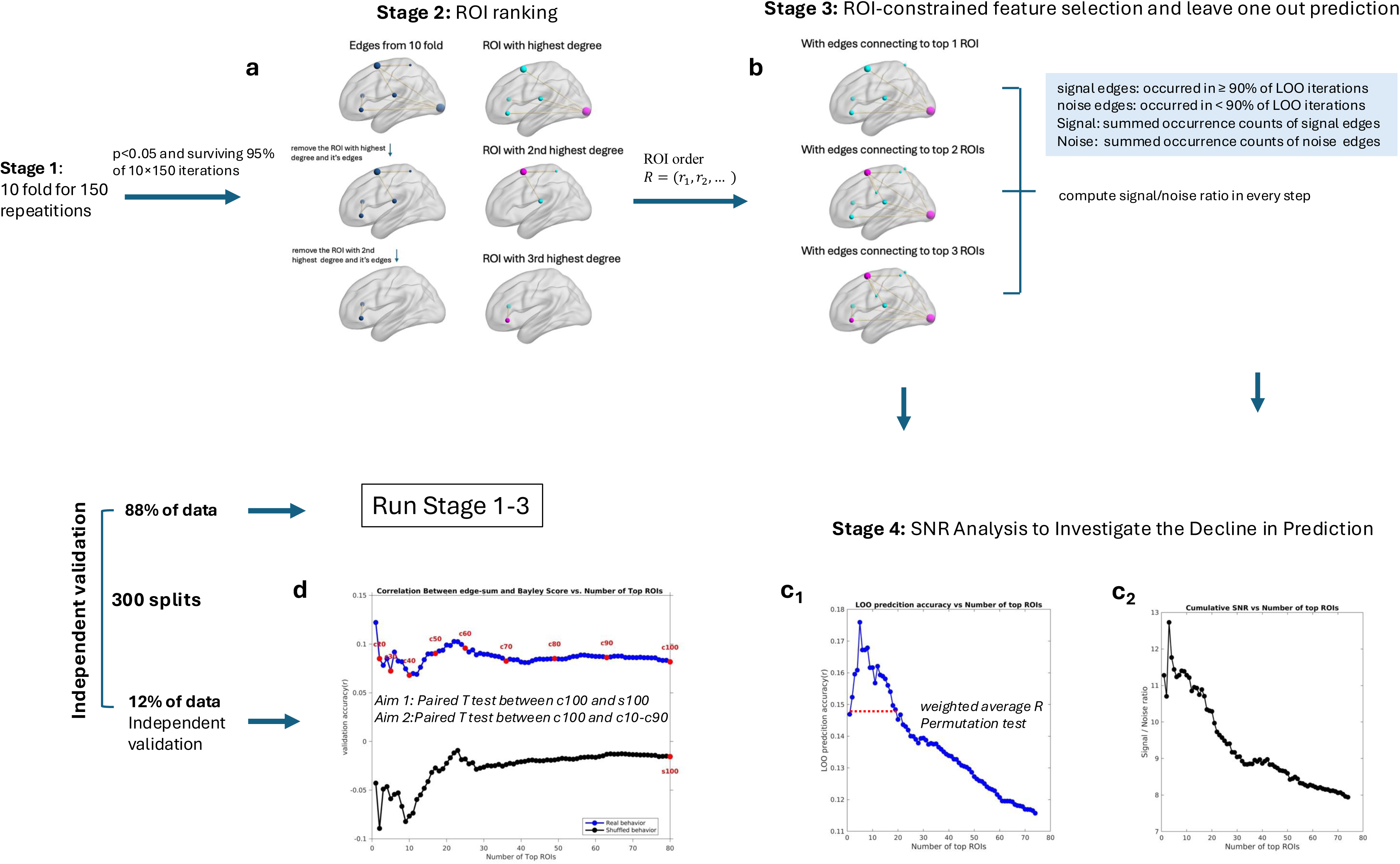
ROI-constrained CPM pipeline with ROI-constrained feature selection and independent validation. (a) Stage-2 — ROI ranking: ROIs were sorted by iteratively removing the highest-degree ROI to define a high-to-low ROI degree order. No Stage-1 edges were carried forward as features; only the ROI order *R* was passed to Stage 3. (b) Stage-3 — ROI-constrained CPM: For each ROI-set size (*n* = 1…|R|), edges that survived p<0.05 and connected to at least one of the top-n ROIs were selected via LOO to calculate prediction accuracy. SNR was also computed for each prefix n by dividing the summed occurrence counts of signal edges (occurred in ≥ 90% of LOO iterations) by those of noise edges (occurred in ≥ 90% of LOO iterations);(c) Stage-4 — SNR analysis: As progressively lower-degree ROIs were added, SNR steadily decreased(c_2_), mirroring the decline in predictive accuracy(c_1_). And SNR and prediction accuracy were strongly correlated (r = 0.96, p < 0.0001). We conducted a permutation test to confirm that the observed weighted average prediction performance (weighted by degree of each ROI) across all ROI-set sizes n exceeded chance; (d) Independent validation: For each ROI-set size (n = 1…|R|), validation subjects’ network strength—sums of signal edges incident on the top-n ROIs—were correlated with behavior to yield *r*(n) curves. The cumulative distribution of edge counts across ROIs was computed, and thresholds were set at 10 %, 20 %, …, 100 % coverage of total edges. Aim 1 — Check whether all signal edges contribute predictive information: Paired T test between observed and shuffled behavior at coverage 100%. Aim 2 — Check whether the predictive effect declines as signal edges of low degree ROIs are added: Paired t-tests compared performance at each intermediate coverage level (10–90 %) with that at 100 %. Abbreviations: c10, coverage 10%; s100, shuffled behavior at coverage 100%; ROI, region of interest; LOO, leave one out; SNR, signal to noise ratio.

**Figure 1.**
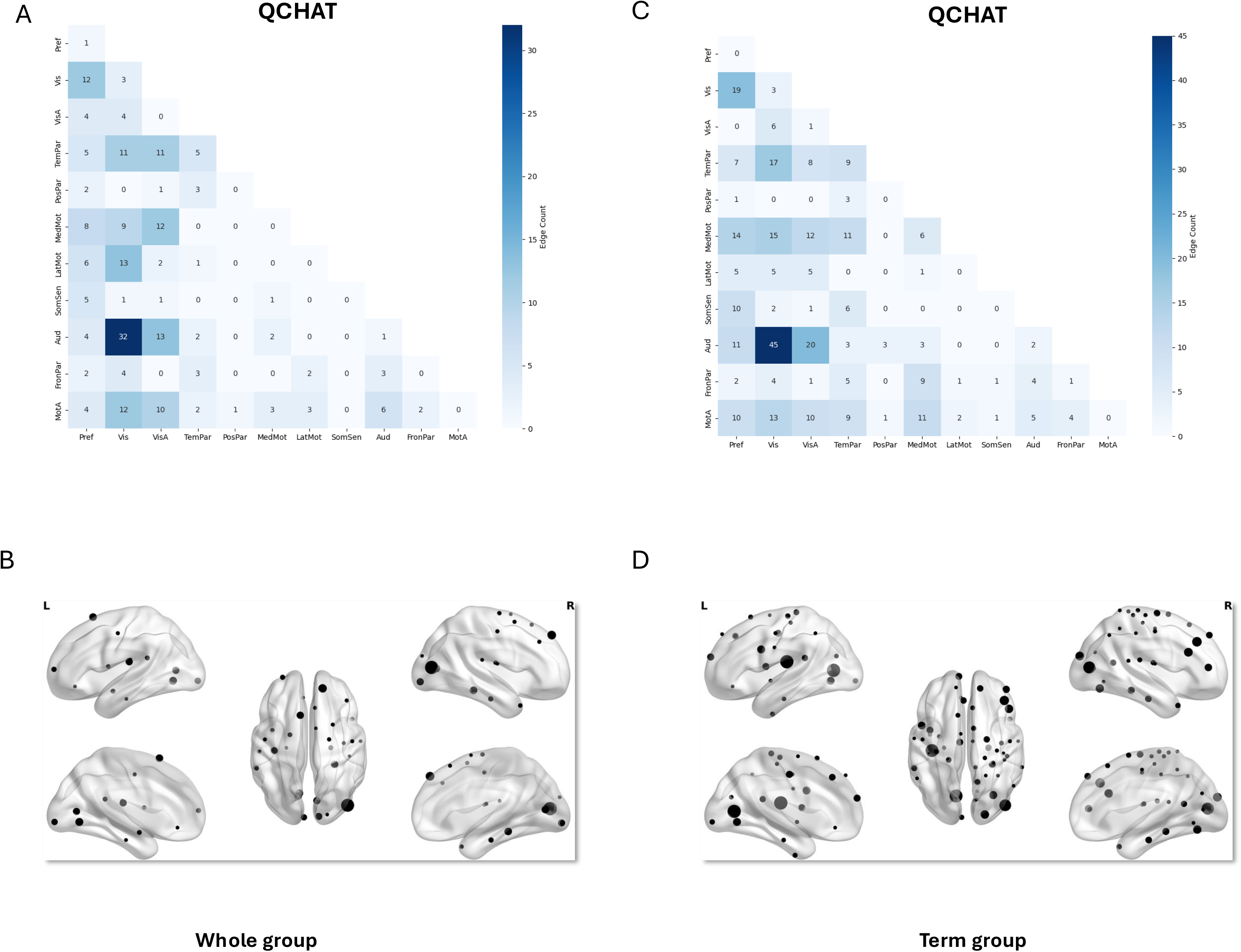
Negative networks: QCHAT predictive anatomy(term cohort). (A,C) Connections plotted as the number of edges within and between each pair of canonical networks for the whole cohort and term cohort respectively(left to right). (B,D) High-degree ROIs for the same models. Only ROIs with a degree ≥ one-sixth of the highest ROI are displayed; node size is proportional to degree.

Stage 3 — ROI-constrained feature selection and leave-one-out (LOO) prediction: We adopted a ROI-constrained variant of CPM. For each prefix size *n* = 1,…,|R| (looping over the ordered ROIs) and each sign, we performed LOO: (1) Feature selection: Within each LOO training set (excluding the held-out subject), we computed partial correlation between each edge and behaviour(the same covariates applied), retaining edges meeting p<0.05 that connected to at least one of the top-n ROIs (Figure 1b). (2) Summary edges: For each subject computed *S*^+^ (sum of positive selected edges) and *S*^-^(sum of negative selected edges). (3) Prediction: We fitted separate linear models on the LOO training set for *S*^+^and *S*^-^ and predicted behaviour of the left-out subject. (4) Edge stability across LOO: After LOO completed for a given n, retaining edges occurring in ≥90% of LOO iterations as *E^+^_final_* and *E^−^_final_(n)*. Stage 3 yielded, for each *n*, sign-specific LOO accuracy (Figure 1c1) and training-stable edge sets. As figure 1c1 shows, as the number of top ROIs increases, the prediction accuracy goes down.

Stage 4 —Signal to noise ratio (SNR) analysis to investigate the decline in prediction: To investigate why the predictive accuracy decreased as more ROIs were added, we examined how the inclusion of lower-degree ROIs affected the SNR.

For each prefix size *n* = 1,…,|R| (looping over the ordered ROIs), we recorded occurrence of all edges across the LOO iterations in Stage 3. These edges were divided into two categories based on a stability threshold of 0.9 × (number of LOO iterations): (1) Signal edges: edges with occurrence counts ≥ threshold; (2) Noise edges: edges with occurrence counts < threshold.

The SNR for each prefix *n* was then computed as

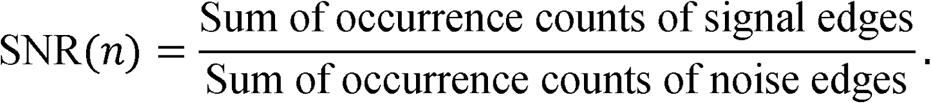

As progressively lower-degree ROIs were added in the whole-cohort model of composite cognition, the SNR consistently declined (Figure 1c2). And SNR was strongly correlated with LOO prediction accuracy (r = 0.96, p < 0.0001; Figures 1c1–1c2). Similar SNR–prediction-accuracy relationships were observed across all behaviours and cohorts, indicating that the decrease in prediction arises from a reduced SNR as low degree ROIs are incorporated.

### Permutation testing of prediction accuracy

After demonstrating that the decline in predictive accuracy with increasing number of top ROIs reflects a reduction in SNR, we conducted a permutation test to confirm that the observed weighted average prediction performance (weighted by degree of each ROI) across all ROI-set sizes *n* (figure 1c1) exceeded chance. Across Stages 1–3, behavioural labels were randomly permuted while preserving the full analytical pipeline. Because QCHAT scores in this healthy neonatal cohort included many tied values, we excluded permutations in which >40% of the scores were reassigned to their original positions. In cases where no edges survived across the 10 folds in Stage 1, the corresponding prediction performance was assigned a value of zero. For each permutation, we computed the weighted average R across all ROI-set sizes *n*. The one-sided *p*-value was calculated as following:

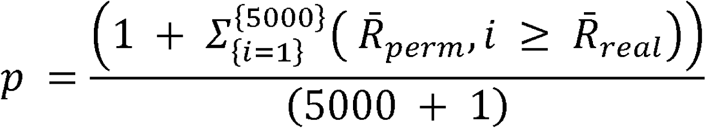

5000= total number of permutations. *R_perm,i_* = weighted average correlation between predicted and permuted behavioral scores across all ROI set sizes *n*. *R_real_* = weighted average correlation between predicted and observed behavioral scores across all ROI set sizes *n*.

### Independent validation of signal edge effects and ROI-dependent decline

We had previously established that the reduction in predictive accuracy is driven by the growing contribution of noise edges originating from low-degree ROIs. It remains unclear whether all signal edges—including those connected to low-degree ROIs—contribute predictive information, and whether prediction accuracy decreases as signal edges from low-degree ROIs are added.

In order to investigate this, we conducted an independent validation analysis. Approximately 88% of the data were used for running stage 1-3, while the remaining 12% were reserved for independent validation. The split was repeated for 300 times. All feature discovery and model fitting used training data only; validation data were untouched until final testing.

To preserve the discretized outcome distribution, we stratified the split by exact QCHAT score value: for values with ≥8 subjects we assigned ∼88% to training and ∼12% to validation; values with <8 subjects were pooled before applying the same proportion.

For each sign and each prefix size *n = 1,…,|R|* (looping over the ordered ROIs), we used the training-stable edge sets *E^+^_final_* (signal edges from Stage 3): (1)For each validation subject, we computed *S*^+^/*S*^-^by summing all edges incident on the top-*n* ROIs; (2)Each subject’s s^±^ was correlated (Pearson’s *r*) with behaviour to yield predictive-accuracy curves *r^±^(n)*; (3) A single-shuffle was performed by permuting the behaviour once per split to estimate chance-level *r^±^(n)*values (Figure 1d).

Aim 1 — Do all signal edges contribute predictive information? To evaluate whether the entire set of signal edges retains predictive power, we focused on the largest ROI set (*n = |R|*) and did paired *t*-tests(fisher-z transformed r value) between real behaviour and shuffled behaviour across 300 splits to test whether the final model’s correlation remained significantly above the single shuffle (Figure 1d).

Aim 2 — Does the predictive effect decline as signal edges of low degree ROIs are added? To examine how expanding the ROI set influences performance, we quantified accuracy as a function of cumulative edge coverage. For each behavior, the cumulative distribution of edge counts across ROIs was computed, and thresholds were set at 10 %, 20 %, …, 100 % of total edges. For each coverage level, the corresponding predictive correlations *r(n)* were extracted. Across 300 splits, paired *t*-tests (Fisher-z transformed r value) compared performance at each intermediate coverage level (10–90 %) with that at 100 % (Figure 1d).

### Statistical evaluation and reporting

The following two statistical test significance will be reported: (1) permutation test of average *R* across all ROI-set sizes *n*; (2) independent validation test on the largest ROI set (*n = |R|*). Pooled mean validation accuracy (r□) at *n = R* with 95% confidence interval will be reported.

Significance versus the behaviour-shuffle null at *n= R* is tested with a paired t-test across splits; we report t(df), exact p, and Cohen’s *d_Z_*(see Table 2). Correlations were Fisher-z transformed for inference and back-transformed for presentation.

**Table 2.** Statistics of permutation and independent validation.

|  | Permutation |  | Independent validation |  |  |  |  |  |
| --- | --- | --- | --- | --- | --- | --- | --- | --- |
| | weighted average R<sup>a</sup> | p<sup>a</sup> | $\bar{r}_{\text{validation}}^b$ | 95% CI<sup>b</sup> | $\bar{r}_{\text{shuffled}}^b$ | t(299)<sup>b</sup> | p<sup>b</sup> | Cohen's $d_z^b$ |
| whole cohort (n=397) |  |  |  |  |  |  |  |  |
| Q-CHAT | 0.10 | 0.052 | -0.09 | -0.10- -0.07 | 0.001 | -9.3 | 2.63E-18 | -0.54 |
| term cohort (n=277) |  |  |  |  |  |  |  |  |
| Q-CHAT | 0.07 | 0.237 | -0.07 | -0.09- -0.05 | -0.003 | -5.5 | 5.74E-8 | -0.32 |

<sup>a</sup>Permutation test: We conducted a permutation test (5000 iterations) to confirm that the observed weighted average prediction performance (weighted by degree of each ROI) across all ROI-set sizes (Figure 1c1) exceeded chance levels. Across Stages 1–3 (Figure 1), behavioral labels were randomly permuted while maintaining the full analytical pipeline. For each permutation, the average correlation R across all ROI-set sizes was computed. A one-sided p-value was then computed as the proportion of permutations whose weighted average correlation exceeded the observed value.
<sup>b</sup>Independent validation: Significance of validation versus the behavior-shuffle null on the largest ROI set ( $n = |R|$ ) is tested with a paired t-test across 300 splits; we report $t(df)$ , exact $p$ , Cohen's $d_z$ , pooled $\bar{r}_{\text{validation}}$ with 95% CI and $\bar{r}_{\text{shuffled}}$ .

We considered two cases. First, when performance at 100% coverage was significantly above chance, we further asked whether 100% coverage also yielded the best validation accuracy compared to other coverages. In most behaviors, the highest validation accuracy was observed at 100% coverage, indicating that predictive performance generally improved as signal edges associated with low-degree ROIs were added. In this case, we reported the signal edges that survived in 90% of splits at 100% coverage. Second, for behaviors in which the best coverage was below 100%, this was typically because a small subset of top ROIs accounted for a disproportionately large share of the total edges. If 100 % coverage is still significant above chance, we still reported the signal edges that survived in 90% of splits at 100% coverage(we compared the connectivity patterns at the best coverage and at 100% coverage and found them to be consistent). If performance at 100% coverage was not significantly above chance, we report the signal edges that survived in 90% of splits at the last coverage level that remained significantly above chance. In both cases, all other tested coverage levels before the coverage we report also showed significant differences between real and shuffled data (paired t-test, p < 0.05).

The connectivity results are presented in a network format. Resting-state networks were defined using group-level independent component analysis of term-born infants scanned at 43.5–44.5 weeks postmenstrual age in the developing Human Connectome Project (dHCP) dataset (Eyre et al., 2021). Each ROI from the parcellation was assigned to the network with the highest spatial overlap (winner-takes-all criterion).

## Results

For Q-CHAT, predictive performance was evaluated with a degree-weighted permutation test and an independent validation against a behaviour-shuffled null (Table 2). In the whole cohort (n = 397), the weighted-average prediction correlation was R = 0.10 (permutation test of average R across all ROI-set sizes p = 0.052), and the pooled validation correlation differed from the shuffled null (r□= −0.09, 95% CI −0.10 to −0.07; shuffled r□= 0.001; paired t(299) = −9.3, p = 2.6 × 10□¹□, Cohen’s dz = −0.54). In the term-born cohort (n = 277), the weighted-average correlation was R = 0.07 (permutation test of average R across all ROI-set sizes p = 0.237; r□= −0.07, 95% CI −0.09 to −0.05; shuffled r□= −0.003; paired t(299) = −5.5, p = 5.7 × 10□□, Cohen’s dz = −0.32). The preterm-only model did not reach significance R = 0.03 (permutation test of average R across all ROI-set sizes p = 0.537). The preterm-only model with the term-equivalent age scanning was not significant R = −0.04 (permutation p = 0.675).

Nodes in the whole cohort model were primarily in right middle occipital gyrus, right superior medial gyrus and left lingual gyrus (Figure 2B), while nodes in the term cohort model were mainly in left calcarine gyrus, left rolandic operculum and right middle occipital gyrus (Figure 2D). In both models, the strongest connection was between visual network and auditory network, as well as between visual association network and auditory network. Medial motor, temporoparietal, and prefrontal networks were also involved (Figure 2A-2C).

## Discussion

This study shows that neonatal resting-state functional connectivity predicts Q-CHAT outcomes at 18 months in the whole cohort and in term-born infants when analyzed with a stability-driven ROI-constrained CPM framework. In both statistically significant models, predictive features were centered on occipital and adjacent cortical regions, and the strongest edges linked visual and visual-association networks with auditory systems. Medial motor, temporoparietal, and prefrontal networks also contributed. Together, these findings suggest that later variation in Q-CHAT score is associated with neonatal large-scale functional organization at birth, with on sensory and multisensory systems being the most strongly statistically associated.

The most prominent result was the consistent involvement of visual cortex and visual-association regions. In both the whole-cohort and term-born models, high-degree nodes were concentrated in occipital areas, and the dominant predictive edges linked visual systems with auditory networks. This pattern is developmentally plausible. Sensory systems are among the earliest large-scale networks to become functionally organized, and early visual development is increasingly implicated in later autism-related phenotypes (Doria et al., 2010; Fransson et al., 2009; Girault et al., 2022).

These findings align with a growing literature suggesting that atypical sensory development is a core component of autism risk. Prospective imaging work has linked infant visual brain development to inherited genetic liability for autism, while broader developmental models argue that autism emerges from early alterations in systems supporting perception, attention, and social engagement (Girault & Piven, 2020; Girault et al., 2022). This link is directly relevant to the present findings: if inherited liability for autism is expressed in the early development of visual brain systems, then neonatal connectivity centred on visual and visual-association cortex may represent an early—and partly heritable—substrate of later autism-related traits reported by Girault et al. (2022), and state explicitly how they converge with the visual-hub pattern observed here. Our results extend this account to the neonatal period by showing that connectivity centered on visual and multisensory pathways is associated with later Q-CHAT variation.

The prominence of visual-auditory and visual-association–auditory coupling is especially notable. Early social communication depends on the integration of visual and auditory information, including facial movements, gaze, and speech-related cues (Michon et al., 2022). Disruption in these pathways could plausibly contribute to later individual differences in Q-CHAT scores. Although Q-CHAT is not a direct language measure, it captures early autism-related dimensions closely linked to social orienting and communication. The present results therefore point to neonatal multisensory organization as a plausible early substrate of later autism-related variation.

The predictive models were not limited to sensory cortices alone. Medial motor, temporoparietal, and prefrontal networks were also involved. This broader architecture suggests that Q-CHAT scores may reflect altered coordination between sensory systems and higher-order associative networks rather than local sensory differences alone. That interpretation is consistent with recent neonatal work showing that dynamic connectivity states at birth are associated with later Q-CHAT-related behaviours (França et al., 2024). Taken together, these findings support a distributed-network account in which early autism-related variation is rooted in the organization of sensory-associative systems.

Prediction was significant in the whole cohort and in the term-born cohort, but not clearly in the preterm-only cohort based on the results provided. This does not imply that preterm-born infants lack relevant neonatal predictors. Preterm birth is associated with widespread alterations in structural and functional connectivity, which may increase heterogeneity in the pathways linking neonatal brain organization to later behavioral traits (Eyre et al., 2021; Rogers et al., 2018). The absence of a clear preterm-only Q-CHAT model may therefore reflect greater heterogeneity, reduced subgroup power, or weaker feature stability rather than lack of neural signal.

Even so, the whole-cohort result indicates that stable predictive features for Q-CHAT can be recovered when analysis prioritizes hub-related and reproducible edges. This highlights the methodological value of ROI-constrained CPM. Standard whole-connectome prediction may be diluted by the inclusion of many weak or unstable edges, especially in neonatal data where motion and developmental variability are substantial challenges (Fitzgibbon et al., 2020). By anchoring prediction to stable high-degree ROIs, the present framework improves interpretability while preserving a whole-brain, data-driven design. This strategy is consistent with evidence that hubs occupy central positions in functional brain networks and serve as important anchors of large-scale integration (Power et al., 2013; van den Heuvel & Sporns, 2013).

The choice of predictive framework also warrants comment. CPM is attractive because it is simple, computationally inexpensive, and highly interpretable: the features are individual connections, the model is a linear fit to summary edge strength, and the resulting anatomy can be read directly in terms of networks and hubs (Shen et al., 2017). This transparency contrasts with more flexible but less interpretable approaches such as kernel ridge regression, support vector regression, elastic-net or LASSO regression, and graph- or convolutional-neural-network models (e.g., BrainNetCNN), which can capture nonlinear and multivariate structure but typically require larger samples, are more prone to overfitting, and yield weights that are harder to map onto specific circuits. The ROI-constrained variant used here preserves CPM’s interpretability while mitigating one of its main weaknesses—sensitivity to large numbers of weak or unstable edges—by anchoring feature selection to reproducible high-degree ROIs, which improved stability in these noisy neonatal data. Its disadvantages are that it retains CPM’s assumptions of linearity and mass-univariate feature selection, that results depend on the parcellation and hub-definition choices, and that restricting features to hubs could in principle discard predictive signal carried by peripheral edges; the independent-validation analysis was included partly to test whether broadening the ROI set systematically improved prediction.

Our hub ranking relied on iterative removal of the highest-degree node from the sign-specific stable graph. Degree is the simplest centrality measure and is well suited to identifying the most densely connected regions, but it is not the only option. Alternative measures—betweenness, eigenvector centrality, PageRank, or participation coefficient—capture different aspects of network topology, such as bridging between modules or connection to other highly connected nodes, and could select a somewhat different set of anchor ROIs. Because the predictive features are ultimately the edges incident on the selected ROIs rather than the ranking metric itself, and because the stable graph was dominated by a small number of strongly connected occipital hubs, we expect the broad anatomy to be robust to the specific measure used; we did not, however, compare these measures exhaustively, and a systematic evaluation of alternative hub-selection criteria—and of fully data-driven approaches that bypass hub selection altogether—is a useful direction for future work.

The present findings also complement recent work linking neonatal connectivity to later Q-CHAT score. França et al. showed that neonatal dynamic functional connectivity is associated with atypical social, sensory, and repetitive behaviours measured by Q-CHAT at 18 months (França et al., 2024). The two studies converge in implicating neonatal functional connectivity and sensory systems in the early expression of autism-related traits, but they differ in approach and emphasis: França et al. (2024) characterised time-varying (dynamic) connectivity using an association-based analysis, whereas the present study models static whole-brain connectivity within an out-of-sample predictive framework and localises the predictive signal more specifically to visual and audiovisual pathways. Our results extend that observation by showing that static whole-brain connectivity, analyzed within a stability-driven predictive framework, also carries meaningful information about later Q-CHAT variation. Importantly, the anatomy identified here points specifically to visual and audiovisual pathways as central components of that signal.

Several limitations should be noted. First, the Q-CHAT results are summarized briefly, and interpretation would be strengthened by fuller reporting of effect sizes, validation statistics, and subgroup-specific details. Second, Q-CHAT indexes continuous autism-related traits rather than clinical diagnosis, so the present models should be interpreted as predictors of trait variation within a healthy population rather than diagnostic biomarkers. Because this cohort is unselected and only a small proportion of infants would be expected to receive a later autism diagnosis—and because Q-CHAT scores here fall largely within the typical range—the present findings should be read as describing normative variation in autism-related traits and the neural systems underlying it, and should not be assumed to generalise to clinical autism without validation in enriched-risk or diagnosed samples. Third, rs-fMRI alone cannot determine whether the observed predictive features reflect altered social attention, multisensory integration, maturational timing, or a combination of these processes.

Future work should test whether these visual-auditory predictors generalize to later autism diagnosis, social communication measures, or longitudinal developmental trajectories. It will also be important to determine whether preterm-born infants show distinct Q-CHAT-related predictive architectures that require larger samples or alternative analytic strategies to detect. More broadly, combining neonatal rs-fMRI with diffusion imaging, structural MRI, and longitudinal behavioral phenotyping may help clarify how early sensory-associative organization contributes to later autism-related outcomes.

In conclusion, neonatal functional connectivity predicts later Q-CHAT variation, and the most stable predictive features are concentrated in visual, visual-association, and auditory systems. The prominence of visual-auditory coupling, together with contributions from temporoparietal, medial motor, and prefrontal networks, suggests that early autism-related traits may be foreshadowed by the organization of multisensory systems at birth. These findings support the view that autism-related variation is already embedded in the neonatal connectome and may be especially linked to sensory network development.

## Acknowledgement

All calculations were performed on the high performance cluster maintained by the Trinity College Research IT unit. The research was supported by funding through a joint scholarship programme of the China Scholarship Council and Trinity College Dublin. Data used in the preparation of this manuscript were obtained from the National Institute of Mental Health (NIMH) Data Archive (NDA). NDA is a collaborative informatics system created by the National Institutes of Health to provide a national resource to support and accelerate research in mental health. Dataset identifier: 10.15154/854d-1c71. This manuscript reflects the views of the authors and may not reflect the opinions or views of the NIH or of the Submitters submitting original data to NDA.

## Author Contributions

Arun L. W. Bokde conceptualized the study, supervised the project and revised manuscript. Mi Zou developed the innovative methodology, performed the data analysis, and wrote and revised the manuscript.

